# High-level ciprofloxacin resistance in *Pseudomonas aeruginosa* is associated with genomic adaptation, collateral antibiotic susceptibility, and physiological trade-offs

**DOI:** 10.64898/2026.07.29.741402

**Authors:** Abhijeet Sahu, Rohit Ruhal

## Abstract

Antimicrobial resistance in *Pseudomonas aeruginosa* has emerged as a major clinical concern. Ciprofloxacin has potent intrinsic activity against *P. aeruginosa*, but resistance to this antibiotic is increasingly reported in clinical settings. In this study, *P. aeruginosa* ATCC 27853 strain was exposed to gradually increasing ciprofloxacin concentrations for 50 passages (∼100 days). The final selected resistant strain (CIP-R) with 256-fold of minimum inhibitory concentration was studied further. Whole-genome sequencing revealed 12 genomic alterations, including known mutations in the quinolone resistance-determining genes *gyrA* (Thr83Ile), *parC* (Thr177Asn), *parE* (Glu459Lys). A duplication mutation in *nfxB* (Tyr153_Gly154dup) was also observed. In addition to these genes, we observed mutations in *pilA, tadB, psdR, NP446_RS32055* (TRAP transporter permease), multiple *dppA3* variants, and a prophage-associated hypothetical gene (*NP446_RS24255*), which have not been reported previously. The *gyrA* Thr83Ile substitution was conserved in 86.38% of ciprofloxacin-resistant clinical isolates retrieved from the NCBI database. Resistance acquisition was accompanied by slower growth, impaired swimming and swarming motility, diminished surface attachment, reduced biofilm formation. The resistant strain has enhanced β-lactamase activity, and resistance to levofloxacin, cefepime, and meropenem together with sensitivity to piperacillin–tazobactam and aztreonam. This study highlights the *gyrA* Thr83Ile mutation as a key genomic marker for molecular screening of ciprofloxacin resistance and reveals secondary adaptive trade-offs that can be targeted for clinical diagnostic and therapeutic decision-making. In conclusion, achieving high-level ciprofloxacin resistance in *P. aeruginosa* involves non-target-site genomic adaptations and physiological trade-offs beyond classical target mutations.

## 1. Introduction

The rapid increase of antimicrobial resistance (AMR) has turned into a huge international public health problem, and which seriously compromises the efficacy of current therapeutic strategies [1]. It is now commonly seen as a serious threat, not only to healthcare systems, but also to the wider areas of society and economic development. According to the current estimations from 2019 at global level, bacterial AMR was involved in over 4.95 million deaths worldwide and was directly accountable for 1.27 million deaths. The above data demonstrate the serious impact of AMR, for which there is an urgent need for further research and reliable treatment methods [2].

*Pseudomonas aeruginosa* is one of the most challenging clinical pathogens due to its remarkable genomic plasticity, metabolic versatility, and extensive intrinsic and acquired resistance mechanisms [3]. *P. aeruginosa* is one of the ESKAPE pathogens and is associated with a variety of healthcare-associated infections such as pneumonia caused by ventilators, bloodstream infections, persistent respiratory infections, burn wound infections, and urinary tract infections in immunocompromised individuals [4,5]. *P. aeruginosa* alone is responsible to cause more than 300,000 AMR-associated deaths each year. According to World Health Organization (WHO) this bacterial species has been classified as high-priority pathogen due to its clinical significance and rapidly increasing resistance to available therapeutics, thus leading to the rapid development of novel antibiotic strategies [6,7].

Ciprofloxacin is one of the major broad-spectrum fluoroquinolone antibiotics that is efficient towards both Gram-negative and Gram-positive bacteria and is utilized in the management of urinary tract, respiratory tract, bone and joint and intra-abdominal infections [8,9]. Ciprofloxacin halts the bacterial growth by inhibiting DNA gyrase and topoisomerase IV, which are critical in DNA replication, DNA repair and transcription [10]. The overuse and inappropriate use of ciprofloxacin have hastened the emergence of AMR which has impaired their therapeutic efficacy [11]. The rise of ciprofloxacin resistance has become a global public health concern, particularly among Gram-negative bacteria like *Pseudomonas aeruginosa* [12], *Escherichia coli* [13], *Klebsiella pneumoniae* [14], *Acinetobacter baumannii* [15], and *Salmonella enterica* [16].

Ciprofloxacin resistance is a complex evolutionary process including multiple genetic and physiological modifications [17]. The most common mechanism of resistance is via mutations in *gyrA*, which encodes DNA gyrase, and is often accompanied by secondary mutations in *parC* and less commonly *parE*, which encode subunits of topoisomerase IV, leading to progressively higher levels of fluoroquinolone resistance [18,19]. In *P. aeruginosa* alterations in the negative regulator *nfxB* of the MexCD–OprJ efflux pump result in efflux pump overexpression [20] and are important in multidrug resistance, whereas in other Gram-negative bacteria like *E. coli* and *K. pneumoniae*, ciprofloxacin resistance is more commonly linked to mutations in the AcrAB–TolC efflux regulatory network [21,22]. These mechanisms are usually acquired after an extended period of insufficient exposure to antimicrobial agents, and yet the evolutionary pathways leading to increased levels of resistance are poorly understood, particularly with respect to clinically identified resistance patterns [23].

The primary goal of our study was to investigate the ability of a ciprofloxacin-resistant (CIP-R) strain of *Pseudomonas aeruginosa* to thrive in the presence of high doses of ciprofloxacin under controlled laboratory conditions and to elucidate the mechanisms involved in this adaptation. We further explored the metabolic trade-offs of resistance and the collateral sensitivity patterns that may offer opportunities for alternative therapeutic strategies. Moreover, our results show that long-term exposure to ciprofloxacin allows the bacteria to acquire a level of resistance that allows them to grow and persist without the need to form biofilms.

## 2. Materials and Methods

### 2.1 Bacterial strain, culture conditions and antibiotic preparation

The reference bacterial strain *Pseudomonas aeruginosa* (ATCC 27853) was obtained from the ATCC in the Kwik-Stik format. For experimental work, the culture was revived from a -80 °C glycerol stock. A fresh bacterial culture was aseptically prepared by introducing a tiny aliquot of the frozen microbial stock into 5 mL of sterilized Luria-Bertani (LB) broth (HiMedia, India). The mixture was incubated at 37 °C with 200 rpm shaking for 12–15 hours. The microbial broth was streaked onto the plates made from LB agar (HiMedia, India) using adequate sterilized methods to maintain the purity and viability of the culture. Following incubation, well-isolated colonies were chosen to serve as the initial population for both subsequent phenotypic studies and adaptive laboratory evolution (ALE) tests. The antibiotic Ciprofloxacin (HPLC grade) was bought from Sigma-Aldrich and was utilized as the selective agent during the research. A stock solution (1 mg/mL) of ciprofloxacin was prepared by dissolving it in sterile Milli-Q water. The solution was purified using a 0.22 μm syringe filter (Millex-GP, Merck) for sterility and to eliminate particle pollutants. The purified stock was utilized in all tests requiring antibiotic treatment and kept in suitable storage.

### 2.2 Laboratory generation of ciprofloxacin resistant strain

The MIC of ciprofloxacin (CIP) against *P. aeruginosa* was obtained by broth microdilution method before the start of adaptive laboratory evolution (ALE). CIP was then serially diluted 2-fold on a 96 well plate and administered with 50 µL of a 1:100 adjusted fresh culture. The plate was then examined for bacterial growth by visual inspection of turbidity and determination of optical density at 600 nm after overnight incubation at 37 °C. The MIC was calculated as the lowest concentration with no visible growth and the minimum OD_600_ equivalent to the sterile control. This value was used as a reference for the design of the ALE experiment. Adaptive laboratory evolution (ALE) [24] was initiated at sub-inhibitory concentration corresponding to MIC/6 (0.083 µg/mL) and a parallel control without exposure to antibiotics. An isolated colony was cultivated overnight in LB broth, and the culture density was adjusted to an optical density of 0.08–0.12 using MHB. Subsequently, 50 µL of this microbe was administered into 5 mL of MHB containing an appropriate dose of CIP, with three biological replicates established for both the treatment and control groups. Cultures were maintained at 37 °C with agitation at 200 rpm for 48 hours. After each growth cycle, 1% of the culture (50 µL) was added into the fresh medium with a gradually increasing concentration of CIP to maintain selection pressure. This process was continued for 50 passages, during which the CIP concentration increased up to 11.5 µg/mL.

### 2.3 Minimum inhibitory concentration (MIC) of Parent and CIP-R strains

The MIC of ciprofloxacin against the parent strain (PA) and the ciprofloxacin-resistant (CIP-R) strain was evaluated using broth microdilution with minor changes to established methods [25]. To create an overnight bacterial culture an isolated colony was introduced into 10 mL of sterile MHB and cultivated at 37 °C with shaking for 12–15 hours. The culture was further calibrated to an OD₆₀₀ of 0.12 (about 1–2 × 10⁸ CFU/mL) and further diluted 1:100 to get a workable inoculum of approximately 1–2 × 10⁶ CFU/mL. Ciprofloxacin was diluted two-fold from 256 µg/mL to 0.25 µg/mL in a sterile 96-well microtiter plate by sequentially transferring equal quantities from a stock solution. 100 µL of bacterial suspension was introduced into each well of MHB. The plate was cultured at 37 °C for 18 to 20 hours. The wells were visually assessed for turbidity post-incubation, and the MIC was defined as the lowest concentration at which no apparent growth occurred. Visual observations were validated by measuring OD_600_ using a microplate spectrophotometer (Agilent BioTek). In addition, 50 µL of the wells with no visible growth was plated on LB agar plates and incubated at 37 °C for 18-24 hrs to detect for viable cells. Colony counts were carried out to check the absence of viable bacteria and to confirm the accuracy of the MIC determination for both strains.

### 2.4 Minimum bactericidal concentration (MBC) of Parent and CIP-R strains

The MBC of ciprofloxacin against the parental (PA) and the ciprofloxacin-resistant (CIP-R) strains was determined by the modified broth microdilution technique. One isolated colony was cultivated overnight in MHB at 37 °C with shaking and diluted 1:100 in fresh MHB. Serial two-fold dilutions of ciprofloxacin (256 µg/mL to 0.25 µg/mL) were done in a sterile 96-well plate and the diluted bacterial sample was added to each well. The 96-well plate was incubated at 37 °C for 18-20 hrs and the samples were further processed for the evaluation of bactericidal activity. 3 µL of each concentration well was carefully spotted on LB agar plates after incubation for the parental and CIP-R strains. The spots were then allowed to be soaked in and air dried under aseptic conditions for a short while. Bacterial survival and bactericidal activity of ciprofloxacin was determined by maintaining the plates at 37 °C for 15-18 hours.

### 2.5 Growth kinetics of the Parent (PA) and CIP-R strains

Overnight cultures of the parental (PA) and ciprofloxacin-resistant (CIP-R) strains were separately cultivated in 10 mL LB broth administered from a single well-isolated colony and incubated at 37 °C with agitation at 200 rpm for 12–15 h for growth curve characterization. The microbial cultures were then diluted to OD600 of 0.05 in fresh LB broth. 50 µL of each diluted culture was added to 50 mL of freshly prepared LB broth in Erlenmeyer flasks and incubated at 37 °C with shaking at 200 rpm. Samples were taken at 2 h intervals and the OD_600 was measured with a UV-Visible spectrophotometer (Shimadzu UV-1280) using 1mL of samples. The measurements were carried out up to 24 h when the cultures reached the stationary phase. All the tests were carried out in triplicate for consistency and reliability of the results. Subsequently, development curves were generated by plotting OD_600_ vs time using Microsoft Excel. The specific growth rate (µ) was calculated from the exponential phase of the bacterial growth curve by applying the equation

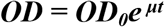

where *OD_0_* is the initial optical density, OD is the optical density at a particular time point t and e is the base of the natural logarithm. The growth rate was estimated in three independent experimental duplicates. The mean and standard deviation were calculated using the obtained numbers using Microsoft excel.

### 2.6 Scanning electron microscopy (SEM) of the Parent (PA) and CIP-R

For scanning electron microscopy (SEM) studies, a single colony of both PA and CIP-R strains were inoculated in 10 mL LB broth and incubated at 37 °C overnight. The culture was diluted to OD600 of 0.4-0.6 with fresh LB broth. A sterile coverslip was put into a petri dish and 100 µL of the diluted culture was carefully spread with a sterile 200 µL pipette tip on the coverslip to form a thin layer. The sample was settled at RT for 20-30 mins and then dried at 37 °C for 3-4 hr. After incubation, the coverslip was washed gently twice in PBS to remove loosely adherent cells. Cells were fixed by adding 200 µL of 2.5% glutaraldehyde and incubating for 30 min at room temperature and overnight fixation at 4 °C. Excess fixative was washed out by three washes in PBS. Samples were dehydrated in an ascending series of ethanol (30%, 50%, 70%, 90% and 100%) for 10 minutes each step [26]. The coverslips were dried at 37 °C for 24 h and then subjected to a vacuum treatment to ensure complete drying. The prepared samples were observed in the last step with a scanning electron microscope (EVO 18, Carl Zeiss).

### 2.7 Motility (Swimming, Swarming and Twitching) determination of Parent (PA) and CIP-R strains

#### 2.7.1 Swimming motility

Swimming motility assay was performed in a slightly modified protocol of that previously described by [27]. Swimming agar plates were prepared using LB medium (10 g/L tryptone, 5 g/L yeast extract and 5 g/L NaCl) containing 0.3% (w/v) agar (HiMedia). A colony of each strain was grown overnight at 37 °C in LB broth. The plates were dried under laminar airflow for about an hour to remove any excess surface moisture before inoculation. A sterile toothpick or a 2 µL pipette micro tip was dipped into the fresh culture and penetrated into the centre of agar surface without touching the bottom of the plate. The plates were then incubated in an upright position at 37 °C for approximately 15 hours to visualise swimming motility.

#### 2.7.2 Swarming motility

The swarming motility assay was performed with minor modifications of a previously published method [28]. Swarming agar plates were prepared by adding 0.5% (w/v) agar (HiMedia) and 5 g/L glucose (HiMedia) to 13 g/L nutrient broth (HiMedia). Prepared plates were kept under laminar air flow for about one hour to remove extra moisture from the surface. Inoculation was performed by spotting 3 µL of microbial culture developed overnight on the middle of the petri plate. The plates were air dried for approximately 15 minutes and incubated at 37 °C for 16-24 hours after inoculation.

#### 2.7.3 Twitching motility

The twitching motility was assay was done as previously described with some modifications [29]. Plates were prepared with LB medium with 1% (w/v) agar (HiMedia) added. A P10 micro tip was used to pick a single colony from the parental or CIP-R strains and gently pushed through the agar to the bottom of the petri dish. The plates were incubated in inverted orientation at 37 °C for 16–24 h. After incubation, the agar medium layer was carefully removed and the bottom of the petri dish was stained with 1% crystal violet for 20 min. The diameter of the twitching zone was measured to determine motility after washing out any residual dye with water.

### 2.8 Biofilm formation assay of the Parent (PA) and CIP-R

The biofilm formation assay was carried out by using a microtiter plate technique with some modifications [30]. One colony of the parental and CIP-R strains was inoculated into 10 mL of LB broth and incubated overnight at 37 °C. The microbial culture was re-suspended 1:100 in fresh LB broth. The diluted culture was added to 100 µL of sterile LB suspension in each well of a 96-well plate. The plate was incubated at 37 °C for 24 h to allow formation of biofilm. After incubation, the culture was aspirated gently and non-adherent cells were removed by washing the wells twice with sterile distilled water. The adhered biofilm was stained by incorporating 125 µL of 0.1% crystal violet reagent to each well and incubated at ambient temperature for 10–15 minutes. The plate was rinsed 2-3 times with water to eliminate excess discolouration and allowed to air dry overnight. The dye had been dissolved by adding 125 µL of 30% acetic acid to each well and incubated at ambient temperature for 10 to 15 minutes. Ultimately, OD_550_ was quantified utilizing a microplate spectrophotometer (Agilent BioTek) to determine biofilm development.

### 2.9 Confocal laser scanning microscopy (CLSM) of the Parent (PA) and CIP-R

Overnight cultures of the parental and CIP-R strain were washed down 1:10 in freshly prepared LB for confocal laser scanning microscopy (CLSM). A sterile glass coverslip was put into a falcon tube so that it remained partly immersed in the medium, to facilitate biofilm growth at the air-liquid interface. Sub-inhibitory concentrations of ciprofloxacin were administered for both bacteria and the setup was incubated for 24–48 hours for biofilm production. Coverslips were gently washed twice with PBS after incubation to remove loosely adhering cells. Then biofilms were dyed with 500 µL of SYTOX® Green dye (5 µM) and incubated in dark for 15–20 min. The extra dye was washed out of the coverslip using PBS. The stained coverslip was positioned on a glass slide (biofilm-side down) and sealed with clear nail polish. Prepared slides were analysed using Olympus FluoView FV3000 confocal microscope. The SYTOX Green dye was stimulated at 504 nm, with fluorescence emission measured at 523 nm.

### 2.10 EPS production of the Parent (PA) and CIP-R

The EPS production ability was evaluated by Congo red agar (CRA) test with minor modifications in the standard procedures [31]. BHI agar plates were prepared by adding 0.08% Congo red dye (HiMedia) and 3.6% sucrose. Fresh parental and CIP-R colonies were streaked on sterile CRA plates. Plates were incubated for 48–72 hrs at 37 °C. Any changes in colony structure and colour were closely monitored after incubation as these traits are indicative of changes in extracellular polymeric substance (EPS) production.

### 2.11 Detection of ß-lactamase enzymes in Parent (PA) and CIP-R

Nitrocefin discs were utilised to detect extended-spectrum β-lactamase (ESBL) activity in *P. aeruginosa*. The parent and CIP-R strains were cultivated overnight and the fresh cultures were diluted to the 0.5 McFarland (∼1.5 x 10⁸ CFU/mL). Then the MHA was cooled to 55 °C and poured onto sterile petri plates containing a sub-inhibitory dose of ciprofloxacin (MIC/4). The bacterial cultures were uniformly spread over agar after solidification using disposable cotton swabs. Plates were kept at 37 °C for 15 hrs. Nitrocefin discs were added to agar plates with sterile forceps and incubated at room temperature for 10–15 min. Colour change in the discs indicates β-lactamase activity.

### 2.12 Comparative antimicrobial susceptibility test (AST) of Parent (PA) & CIP-R

Antimicrobial susceptibility testing (AST) was accomplished with the use of the conventional Kirby-Bauer disk diffusion technique. Overnight parental and CIP-R strain cultures were set to 0.5 McFarland. To create a homogeneous lawn on MHA plates, a disposable cotton swab soaked in the bacterial suspension was used to disseminate the culture throughout the plate. Antibiotic discs of various classes, including ciprofloxacin (5 µg), tobramycin (30 µg), levofloxacin (5 µg), cefepime (30 µg), cefixime (5 µg), ceftazidime (30 µg), fosfomycin (200 µg), polymyxin B (300 units), imipenem (10 µg), meropenem (10 µg), piperacillin– tazobactam (100/10 µg), and aztreonam (30µg) (HiMedia, India)—were carefully placed onto the agar surface under aseptic conditions. At 37 °C plates were cultured for 15–18 hours. The parental and CIP-R strains antibiotic susceptibility patterns were compared by measuring zone of inhibition diameter.

### 2.13 Whole genome sequencing of Parent (PA) & CIP-R

The Illumina sequencing platform with paired-end chemistry (2 × 150 bp) was used to conduct whole genome sequencing (WGS) of the parental and ciprofloxacin-resistant (CIP-R) strains Trimmomatic (v0.39) was used to remove adaptor sequences, low-quality reads and ambiguous bases from raw sequencing reads to create high-quality clean reads. The filtered reads were aligned to the *Pseudomonas aeruginosa* ATCC 27853 reference genome (GCF_024507955.1) using BWA-MEM (v0.7.17). Following alignment, consensus sequences were generated using SAMtools (mpileup). Gene extraction and annotation were performed using BEDTools, along with reference genome annotation files in GFF format obtained from NCBI. Single nucleotide polymorphisms (SNPs) and InDels were identified from mapped data using SAMtools (v0.1.18). BEDTools was then used to further annotate the variants. The variant filtering step was performed using minimum read depth, quality thresholds and other parameters, to ensure reliability. This integrated workflow from raw reads to variant identification enabled us to perform accurate comparative genomic analysis between parental and resistant strains.

## 3. Results

### 3.1 Change in MIC of *Pseudomonas aeruginosa* up to 256-fold under ciprofloxacin pressure

The parent (PA) strain was exposed to adaptive laboratory evolution (ALE) under controlled laboratory circumstances with a progressive, stepwise rise in ciprofloxacin concentration as illustrated in (**Fig 1**). The MIC of ciprofloxacin for the parent strain was 0.5 µg/mL. The parent (PA) strain was repeatedly passaged in MHB with increasing concentrations of ciprofloxacin. We carried out more than 50 passes (∼100 days) from a sub-inhibitory concentration (0.083 µg/mL) up to 11.5 µg/mL. An evolved population (CIP-R) arose at the final dose of 11.5 µg/mL with a MIC of 128 µg/mL of ciprofloxacin. Adaptive laboratory evolution caused *Pseudomonas aeruginosa* to rapidly develop ciprofloxacin resistance. The MIC of the CIP-R strain was 128 µg/mL showing a 256-fold rise over the PA strain (**Fig 2A** and **2B**). Similarly, the minimum bactericidal concentration (MBC) increased from 1 µg/mL in the parent strain to 256 µg/mL in the resistant strain, also representing a 256-fold rise (**Fig 2C**). The resistant strain was passaged on antibiotic-free media for 10 days to test its stability. The strain continues to thrive on ciprofloxacin-containing agar (12 µg/mL) and MIC measurements indicate no resistance decrease (**Fig 2D** and **2E**). This suggests that the acquired resistance was a stable, probable genetic change. These results suggest that *P. aeruginosa* is capable of rapid development and stability of high ciprofloxacin resistance under selection pressure.

**Fig 1.**
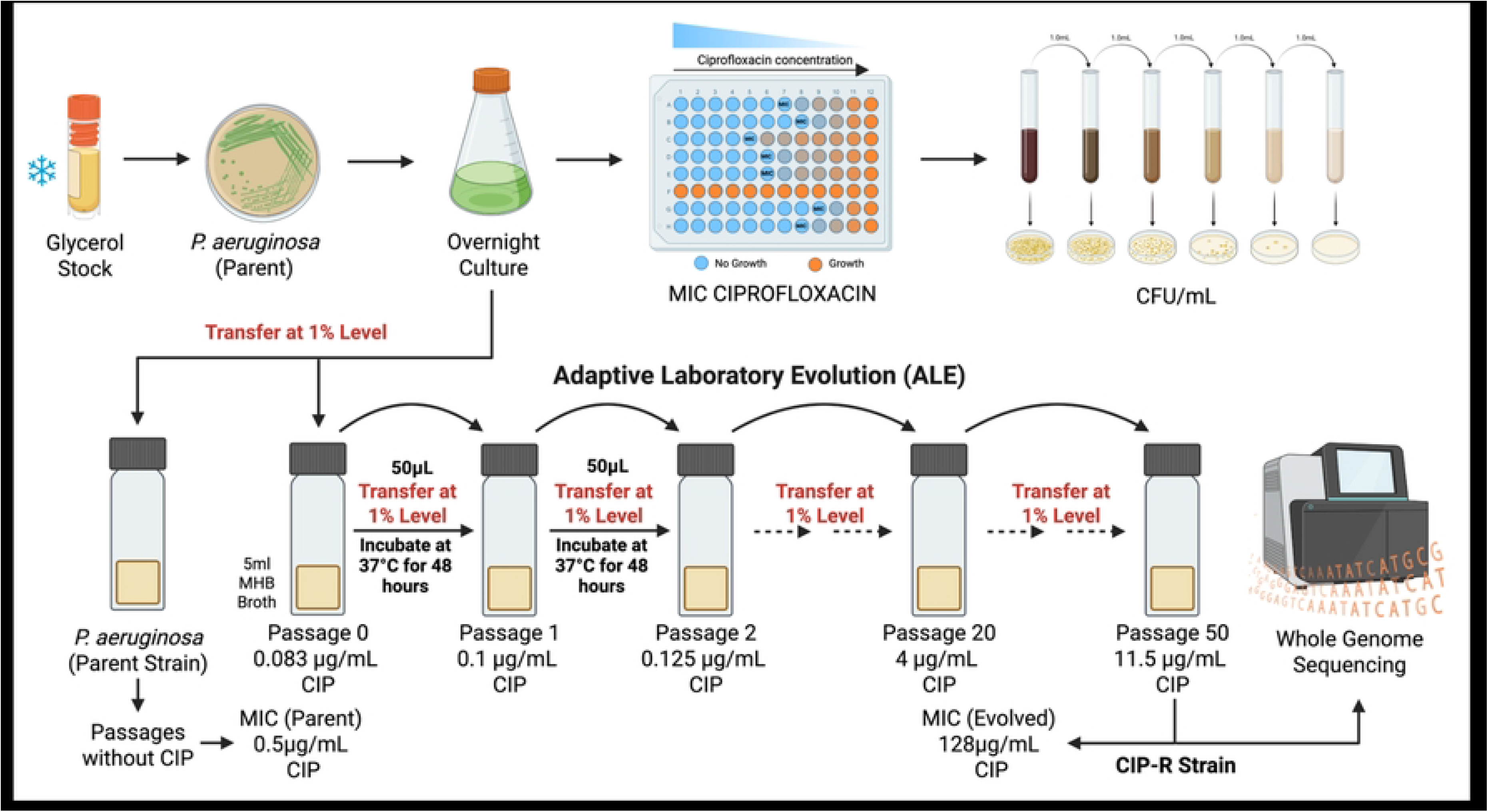
Schematic representation of the adaptive laboratory evolution (ALE) workflow used for the development of a ciprofloxacin-resistant (CIP-R) *Pseudomonas aeruginosa* strain under gradually increasing ciprofloxacin selection pressure. The figure illustrates serial passaging, MIC determination, bacterial enumeration, and whole-genome sequencing analysis performed during the evolution process.

**Fig 2.**
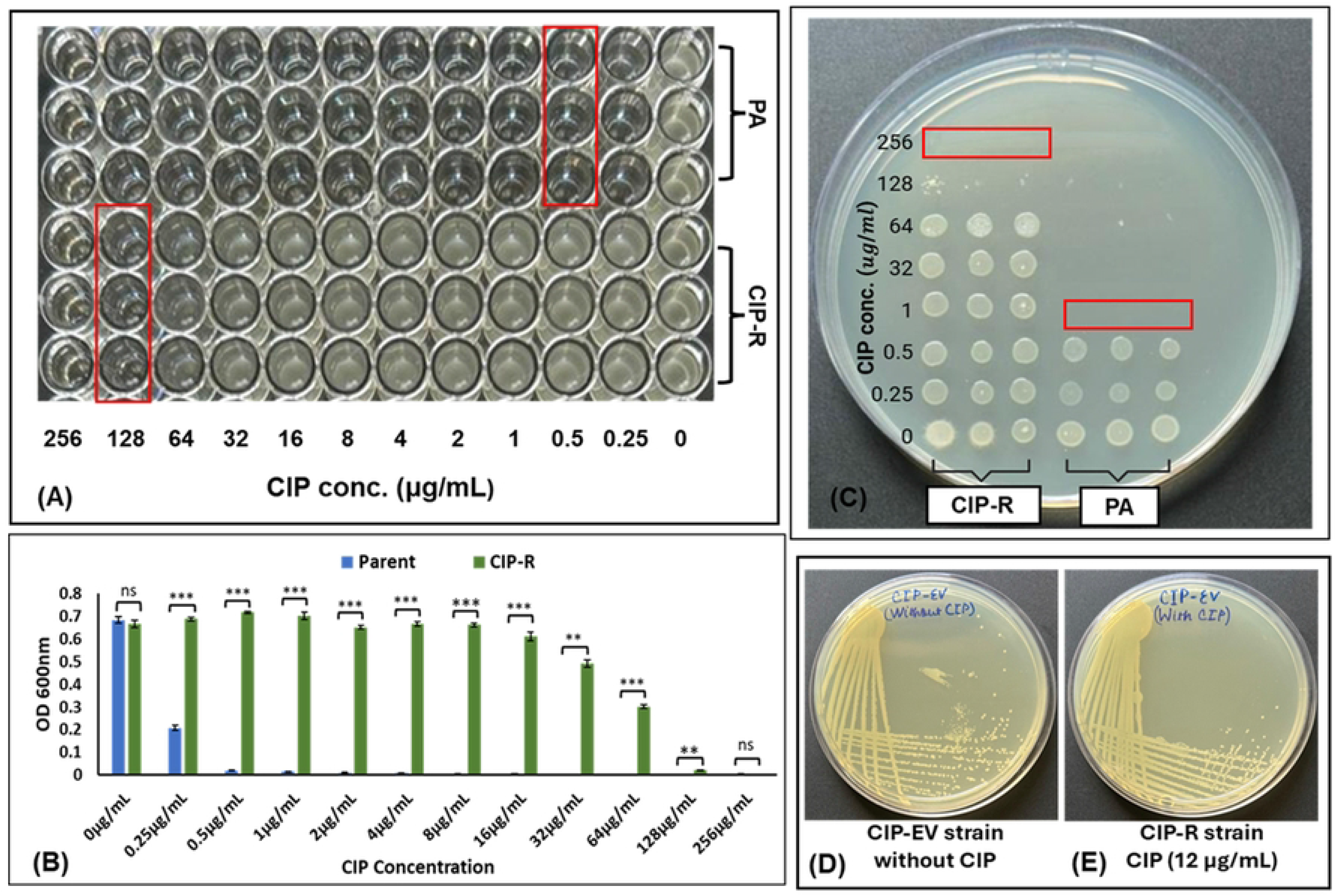
Determination and characterization of ciprofloxacin resistance in the evolved (CIP-R) *Pseudomonas aeruginosa* strain. **(A)** Broth microdilution assay showing MIC values of 0.5 µg/mL for the parental strain (PA) and 128 µg/mL for the CIP-R strain. **(B)** Comparative MIC analysis of PA and CIP-R strains across different ciprofloxacin concentrations with statistical significance analysis. **(C)** Minimum bactericidal concentration (MBC) determination by spot plate assay demonstrating MBC values of 1 µg/mL for PA and 256 µg/mL for CIP-R. **(D, E)** Stability assessment of the evolved strain after ten antibiotic-free passages showing sustained growth of the CIP-R strain in the presence of 12 µg/mL ciprofloxacin.

### 3.2 A significant fitness trade off was observed in ciprofloxacin resistant strain

#### 3.2.1 Altered growth dynamics and fitness cost in the CIP-R strain

Both strains showed clear variations in growth patterns when streaked on LB agar without antibiotic pressure. The colonies of the resistant (CIP-R) strain had a delayed formation on the plates suggesting a decreased growth rate or an increased generation time compared to the parent strain. To study differences both strains were grown in LB broth without ciprofloxacin and growth was monitored for 24 hours measuring OD_600_ at 2-hour intervals. Both strains followed the normal pattern of lag, exponential, and stationary phases, although the two strains did not develop exactly the same. The parent strain began the exponential phase sooner and had a sharper rise in optical density suggesting a faster initial growth. Meanwhile, the CIP-R strain exhibited a delayed start of the exponential growth and a slower rise in cell density throughout the early phases. The resistant strain eventually converged to the parent strain and attained a similar OD during stationary phase (**Fig 3A**). This shows that the resistant strain may still be able to attain a comparable final biomass in non-selective environments, despite a slower early growth. These results were confirmed by the growth rate (µ) as defined in the Materials and Methods section. The growth rate of CIP-R strain was lower (0.783 (± 0.004) h ^-1^) than the parent strain (1.120 (± 0.044) h ^-1^). This variation was statistically significant (P=0.0001) and indicates that the development of ciprofloxacin resistance is associated with a measurable growth fitness cost.

**Fig 3.**
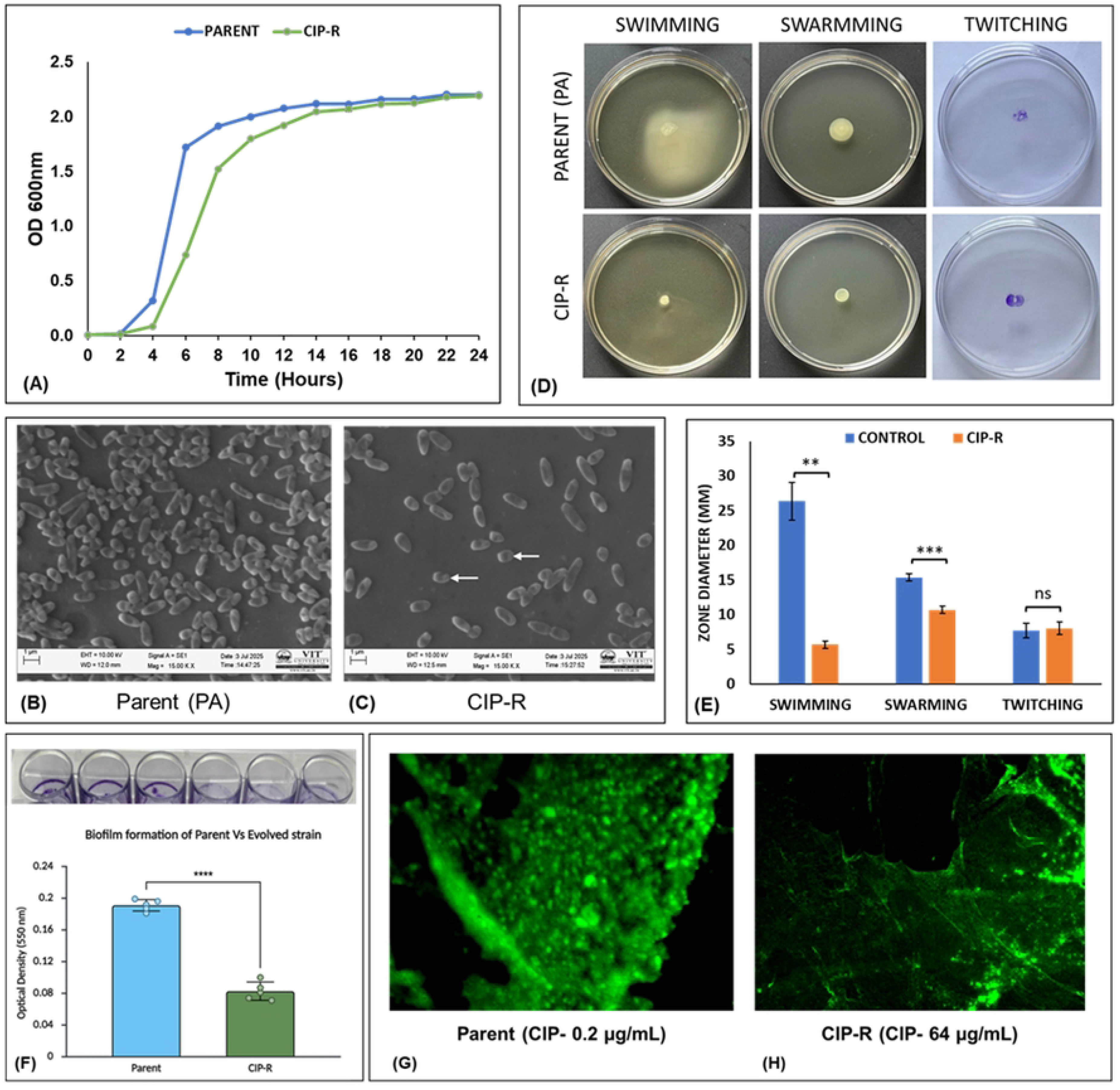
Phenotypic characterization of the ciprofloxacin-resistant (CIP-R) *Pseudomonas aeruginosa* strain in comparison with the parental strain (PA). **(A)** Growth kinetics of PA and CIP-R strains monitored over 24 h by measuring OD at 600 nm at 2 hr intervals. **(B, C)** Scanning electron microscopy revealing morphological differences between the parental and evolved resistant strains. **(D)** Comparative assessment of swimming, swarming, and twitching motility between PA and CIP-R strains, while **(E)** presents the corresponding statistical analysis demonstrating significant reductions in swimming and swarming motility in the CIP-R strain with no significant change in twitching motility. **(F)** Quantitative biofilm formation assay showing reduced biofilm-producing ability in the CIP-R strain relative to the parental isolate. **(G, H)** Confocal laser scanning microscopy images of PA and CIP-R strains exposed to subinhibitory ciprofloxacin concentrations (0.2 µg/mL and 64 µg/mL, respectively), illustrating differences in biofilm architecture and cellular organization.

#### 3.2.2 Scanning electron microscopy reveals reduced surface attachment in the resistant strain

Scanning electron microscopy (SEM) was used to study surface morphology and attachment pattern of the parent (PA) and ciprofloxacin resistant (CIP-R) strains. Both strains were grown on coverslips and imaged using an EVO 18 (Carl Zeiss) microscope at 15,000× magnification. Between the two strains clear differences were observed. The parent strain exhibited tightly packed cells with clear clustering and close surface interaction, suggesting strong attachment and early biofilm formation. The distribution of the CIP-R strain was more dispersed with the cells more evenly distributed and with less aggregation and surface coverage (**Fig 3B** and **3C**). Furthermore, the resistant strain exhibited minor alterations in cell shape and morphology, possibly reflecting changes in physiological conditions. Taken together these results suggest a reduced capacity of the CIP-R strain to adhere and form microcolonies in accordance with its reduced biofilm formation and possible modifications of surface related structures.

#### 3.2.3 Reduced swimming and swarming motility in the CIP-R strain

The motility of the parent (PA) and ciprofloxacin resistant (CIP-R) strains was assessed by evaluating the size of the motility zones for swimming, swarming and twitching under optimum assay conditions. A substantial reduction in motility was observed for the resistant strain in both swimming and swarming **(Fig 3D)**. In swimming motility assays (0.3% LB agar) the average zone diameter of the parent strain was 26.33 ± 2.7 mm and the CIP-R strain had a significantly smaller zone (5.66 ± 0.51 mm, P = 0.0057). Similarly, in swarming assays (0.5% NA agar supplemented with 0.5% glucose), the parent strain showed a zone diameter of 15.33 ± 0.51 mm compared with 10.66 ± 0.51 mm in the CIP-R strain (P = 0.00058).

Results showed statistically significant decrease in swimming and swarming motility after acquisition of ciprofloxacin resistance. On the other hand, twitching motility (1% LB agar) was not significantly different between both strains with similar zone sizes for the parent (7.66 mm) and CIP-R (8 mm) strains **(Fig 3E)**. The apparent fitness trade-offs of the CIP-R strain may have contributed to the decreased swimming and swarming motility but had little effect on twitching motility.

#### 3.2.4 Reduced biofilm formation in the CIP-R strain

Biofilm production of parent strain (PA) and ciprofloxacin resistant strain (CIP-R) was determined by microtiter plate assay and biomass was evaluated by absorbance at 550 nm. The CIP-R strain exhibited a substantial reduction in biofilm formation than the PA strain. As shown in (**Fig 3F**), the OD for the parent strain was 0.17 ± 0.05 whereas that for the resistant strain was significantly lower at 0.081 ± 0.01, showing a reduced capacity to form biofilms in the resistant strain. All studies were performed in triplicate, and findings demonstrate that the CIP-R strain had lower biofilm formation than the parent strain.

#### 3.2.5 CLSM analysis reveals enhanced cell viability in the CIP-R strain

Confocal laser scanning microscopy (CLSM) examined the biofilm architecture and cellular viability of the parent (PA) and ciprofloxacin-resistant (CIP-R) strains cultivated on coverslips under sub-inhibitory concentrations of ciprofloxacin. The parent strain was treated with 0.2 µg/mL and the CIP-R strain was grown in 64 µg/mL and both were stained with SYTOX® Green dye which selectively labels membrane-compromised (dead) cells. Images were acquired using an Olympus FluoView FV3000 microscope. The parent strain formed a dense and well-structured biofilm with high fluorescence signals, suggesting that the proportion of damaged or dead cells in the biofilm was higher (**Fig 3G**). On the contrary, the biofilm of CIP-R strain was less dense and disorganized and the fluorescence intensity decreased at high antibiotic doses, which is consistent with less cell damage and more living cells (**Fig 3H**). These data indicate that the resistant strain has superior cell integrity and hence better tolerance than the parent strain under ciprofloxacin stress.

#### 3.2.6 Slight reduction in exopolysaccharide production in the CIP-R strain

Exopolysaccharide (EPS) production in the parent (PA) and ciprofloxacin-resistant (CIP-R) strains was evaluated using the Congo red agar (CRA) method, followed by quantitative estimation using the phenol–sulphuric acid method. On CRA plates, both strains formed dark, dry colonies, indicating EPS production; however, the parent strain displayed more intense colony characteristics compared to the CIP-R strain, suggesting relatively higher EPS synthesis. This observation was supported by quantitative analysis, where the parent strain produced 175.33 mg/L of EPS, while the CIP-R strain showed a slightly lower value of 165.08 mg/L. Although the difference was modest, it indicates a small reduction in EPS production in the resistant strain. Overall, our data indicate that development of ciprofloxacin resistance is related with a small reduction in EPS synthesis, although the capacity to make extracellular polysaccharides is mainly intact.

### 3.3 Collateral sensitivity and resistance patterns in CIP-R strain

#### 3.3.1 Enhanced β-Lactamase activity in the CIP-R strain

Detection of β-lactamase activity in the parent (PA) and ciprofloxacin-resistant (CIP-R) strains was performed using nitrocefin discs on Mueller–Hinton agar supplemented with sub-inhibitory concentrations of ciprofloxacin (MIC/4). After incubation, nitrocefin discs were placed on the bacterial lawn to assess enzyme activity based on colour change. The parent strain showed no change in the disc colour, indicating the absence of measurable β-lactamase activity under the examined circumstances. In contrast, the CIP-R strain showed a distinct colour change of the nitrocefin disc from yellow to bright red, indicating the presence of active β-lactamase enzymes as demonstrated in (**Fig 4A**). These results imply that the resistant strain has either acquired or up-regulated β-lactamase production, which may be contributing to its altered resistance profile.

**Fig 4.**
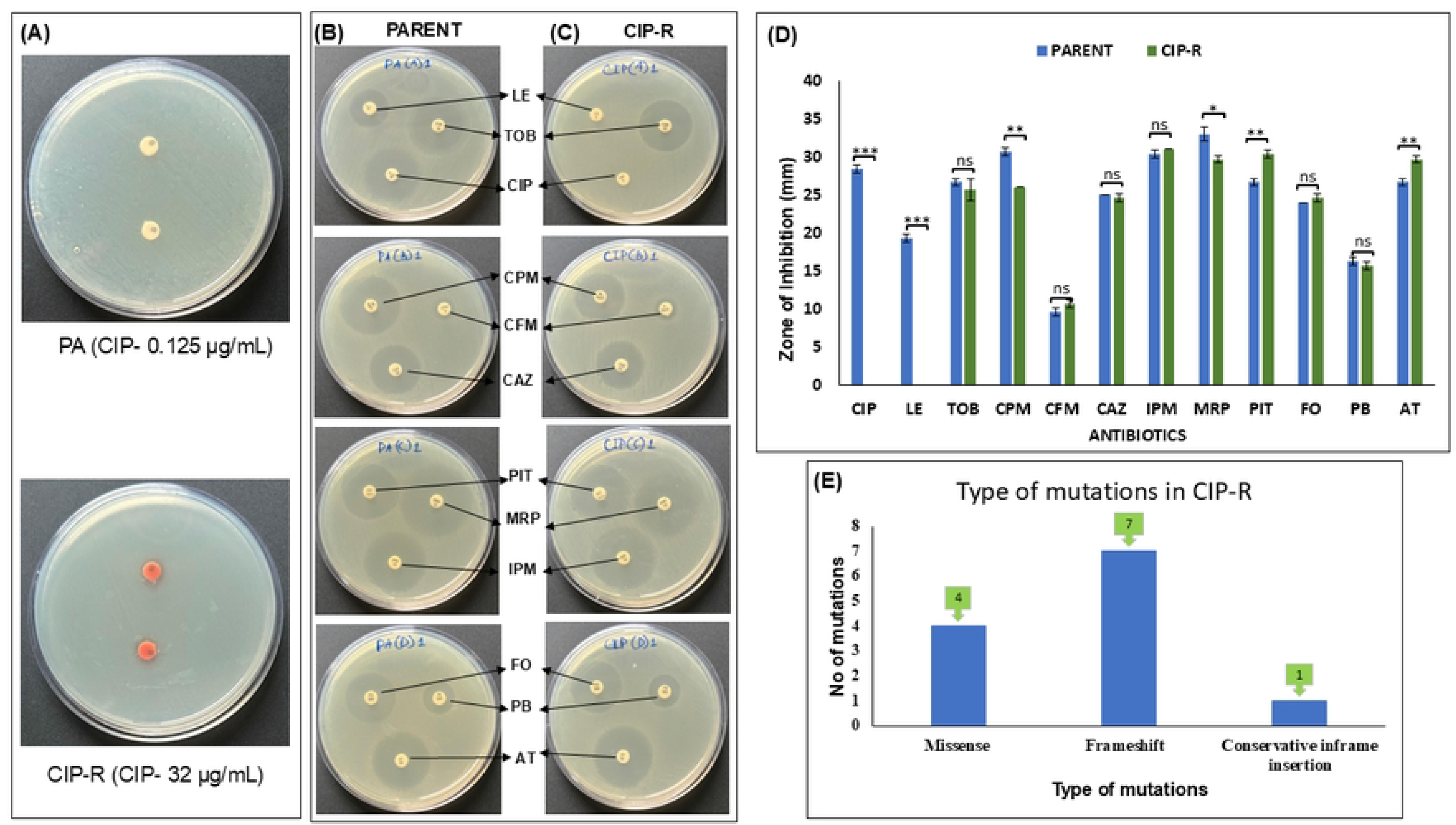
Functional and genomic characterization of the ciprofloxacin-resistant (CIP-R) *Pseudomonas aeruginosa* strain. **(A)** Detection of β-lactamase activity under subinhibitory ciprofloxacin exposure (MIC/4), demonstrating enhanced β-lactamase production in the CIP-R strain, whereas no detectable activity was observed in the parental strain (PA). **(B, C)** Comparative antimicrobial susceptibility profiles of PA and CIP-R strains against multiple antibiotics. **(D)** Statistical analysis of antimicrobial susceptibility patterns highlighting collateral resistance and collateral sensitivity associated with ciprofloxacin adaptation. **(E)** Distribution and frequency of mutation types identified in the CIP-R strain following whole-genome sequencing analysis.

#### 3.3.2 Altered antimicrobial susceptibility profiles following ciprofloxacin resistance evolution

Antimicrobial susceptibility of parental (PA) and ciprofloxacin resistant (CIP-R) strains was assessed by disk diffusion technique (Kirby–Bauer) with a group of antibiotics belonging to multiple classes. The dimensions of the zones of inhibition were documented and evaluated between the two strains, as seen in (**Fig 4B** and **4C**). The CIP-R strain showed a clear decrease in susceptibility to ciprofloxacin, consistent with its evolved resistance. In addition, decreased zone diameters for levofloxacin, cefepime and meropenem demonstrated collateral resistance toward these antibiotics. The MIC of meropenem escalated from 0.5 µg/mL in the PA strain to 8 µg/mL in the CIP-R strain, indicating a 16-fold increase in resistance. In contrast, the resistant strain exhibited greater sensitivity to piperacillin–tazobactam and aztreonam, with wider zones of inhibition in comparison with the parent strain, indicating collateral sensitivity in (**Fig 4D**). Other antibiotics tested showed no noticeable variance in susceptibility between the two strains, including tobramycin, cefixime, ceftazidime, fosfomycin, polymyxin B, and imipenem. Collectively, these data indicate that varying patterns of antibiotic susceptibility, including collateral resistance and sensitivity, correlate with the development of ciprofloxacin resistance.

### 3.4 Genomic determination of CIP-R strain

Whole genome sequencing (WGS) identified 12 genetic alterations in the ciprofloxacin-resistant (CIP-R) strain relative to the parental *P. aeruginosa* (PA), as shown in (**Table 1**). These comprised four missense mutations, seven frameshift mutations, and one conservative in-frame insertion, as represented in the Circos plot **(Fig 5)**. Many of these changes were found in genes that are directly related to fluoroquinolone resistance. Mutations in *gyrA*, *parC* and *parE* that code for key elements of DNA gyrase and topoisomerase IV serve as the most significant targets of the action of ciprofloxacin. Thr83Ile substitution in *gyrA* is a known resistance-associated mutation that probably reduces the binding efficiency of ciprofloxacin to the target and therefore its ability to inhibit DNA replication. In addition, mutations in *parC* (Thr177Asn) and *parE* (Glu459Lys) may further confer resistance by reducing the generation of lethal DNA strand breaks during replication and segregation. Mutations in regulatory and efflux pathways were also seen, in addition to the target-site mutations. The duplication mutation (Tyr153_Gly154dup) in *nfxB*, which encodes a repressor of the MexCD-OprJ efflux pump, may cause loss or impairment of the regulatory control resulting in overexpression of the efflux system and increased extrusion of the antibiotics. A frameshift mutation in *psdR* (Met111fs) regulating dipeptide uptake indicates an adaptation to the metabolic state of the bacteria under long-term antibiotic stress conditions.

**Fig 5.**
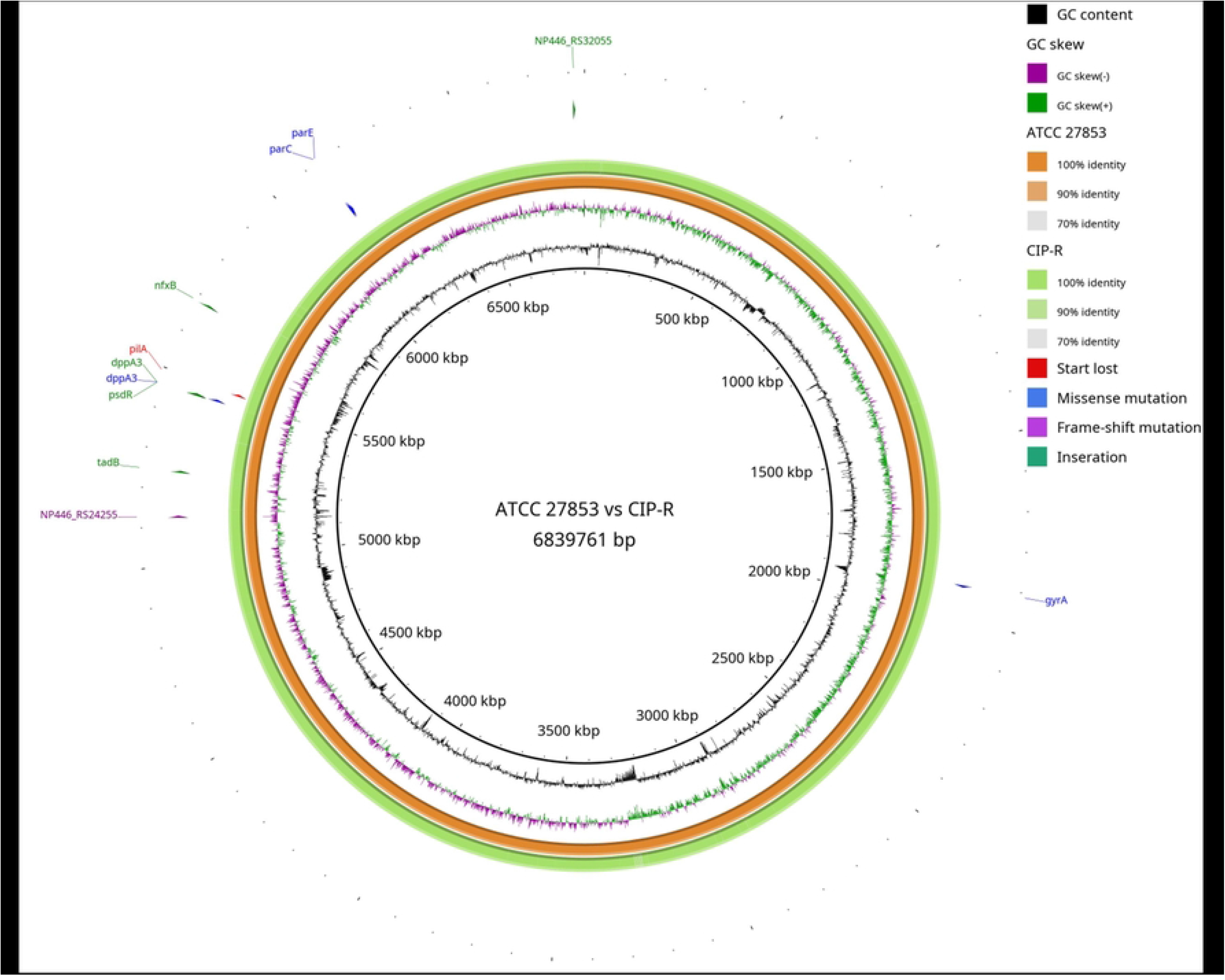
Circos plot showing comparative genomic variations between the parental *Pseudomonas aeruginosa* ATCC 27853 strain and the ciprofloxacin-resistant (CIP-R) strain. The figure highlights the distribution of missense, frameshift, insertion, and start-loss mutations acquired during adaptive laboratory evolution.

**Table 1.** Genomic variants identified in the experimentally evolved ciprofloxacin-resistant *Pseudomonas aeruginosa* strain after prolonged antibiotic selection pressure, with corresponding amino acid changes, mutation types, and predicted gene functions.

| Gene ID | Gene Name | Type of AA Change | Type of Mutation | Description |
| --- | --- | --- | --- | --- |
| NP446_RS29010 | *parC* | Thr177Asn | Missense | DNA topoisomerase IV subunit A |
| NP446_RS29025 | *parE* | Glu459Lys | Missense | DNA topoisomerase IV subunit B |
| NP446_RS09165 | *gyrA* | Thr83Ile | Missense | DNA gyrase subunit A |
| NP446_RS24255 | - | His4fs | Frameshift | Hypothetical protein |
| NP446_RS24785 | *tadB* | Ile213fs | Frameshift | Type 4b pilus Flp biogenesis protein<br>*TadB* |
| NP446_RS25795 | *psdR* | Met111fs | Frameshift | Pseudomonas dipeptide regulator, *PdsR* |
| NP446_RS25800 | *dppA3* | Arg70fs | Frameshift & Missense | ABC transporter substrate-binding protein |
| NP446_RS25800 | *dppA3* | Leu75fs | Frameshift & Missense | ABC transporter substrate-binding protein |
| NP446_RS25800 | *dppA3* | IleGly81ValSer | Missense | ABC transporter substrate-binding protein |
| NP446_RS25935 | *pilA* | Met1fs | Frameshift & Start lost | Type 4 fimbrial precursor *PilA* |
| NP446_RS26965 | *nfxB* | Tyr153_Gly154dup | Conservative inframe insertion | Efflux pump transcriptional repressor<br>*NfxB* |
| NP446_RS32055 | - | Met78fs | Frameshift | TRAP transporter permease |

We also found mutations in virulence and motility genes. Frameshift mutations were found in both *pilA* (Met1fs) and *tadB* (Ile213fs) essential for Type IV pilus formation. Furthermore, several mutations were detected in genes related to transport and metabolic functions. The *dppA3* gene was found to have several mutations including two frameshift mutations (Arg70fs and Leu75fs) and a missense mutation (IleGly81ValSer) which might result in impaired substrate transport. Likewise, a frameshift mutation (Met78fs) in *NP446_RS32055*, encoding a TRAP transporter permease, may affect nutrient uptake processes. These changes together suggest a possible physiological trade-off in which the metabolic efficiency is modified to allow survival in the midst of ciprofloxacin. In addition, a frameshift mutation (His4fs) was identified in the putative protein *NP446_RS24255*. According to PhageScope, this hypothetical gene is in a Caudoviricetes prophage region. Comparative analysis of the ciprofloxacin-resistant evolved strain with previously reported clinical isolates revealed that several mutations occurred within the same resistance-associated genes, although the specific amino acid substitutions differed as shown in (**Table 2**). This trend is consistent with adaptive evolution on shared genetic pathways although the exact mutational alterations may vary in strains adapting to various selective environments.

**Table 2.**
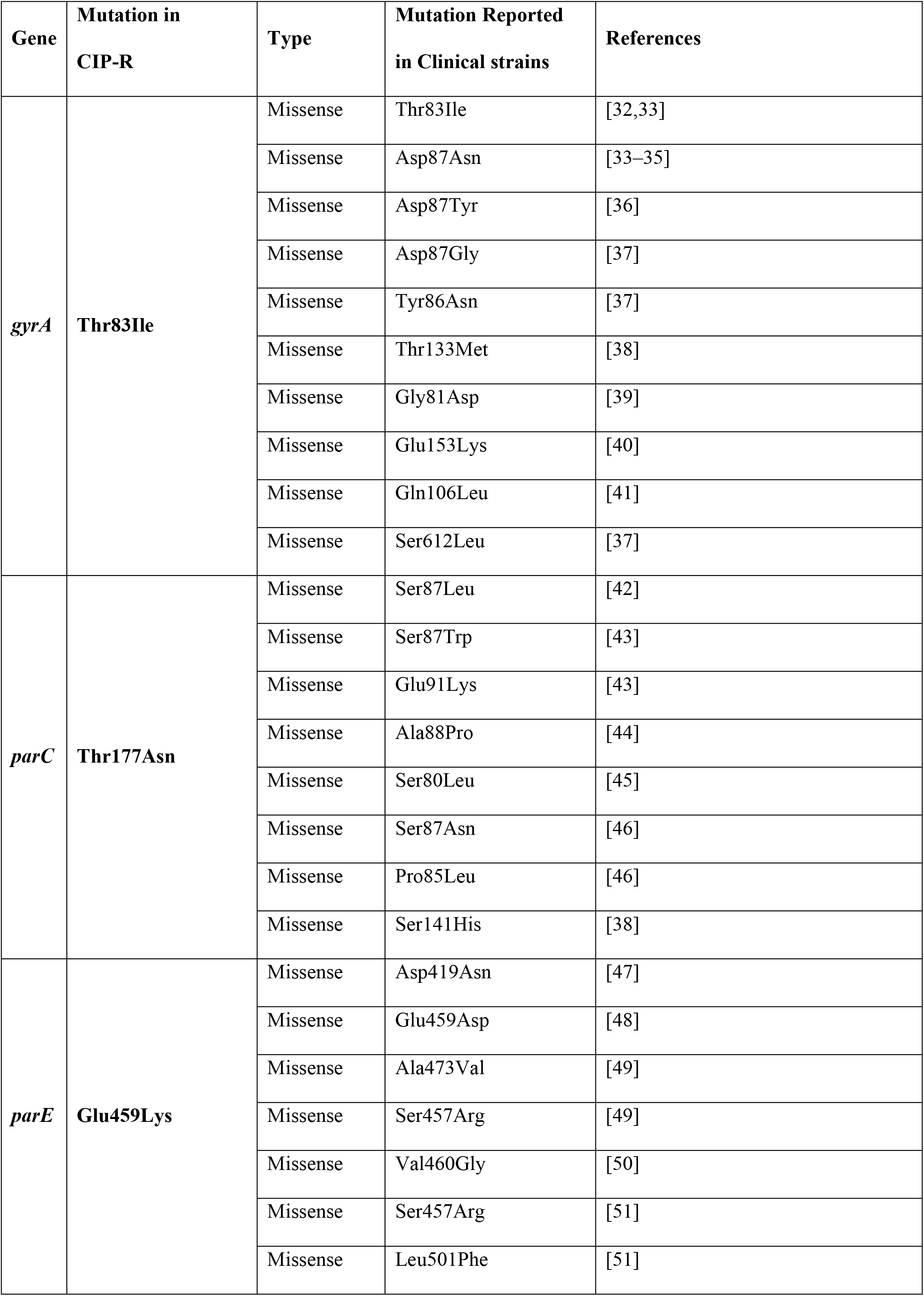
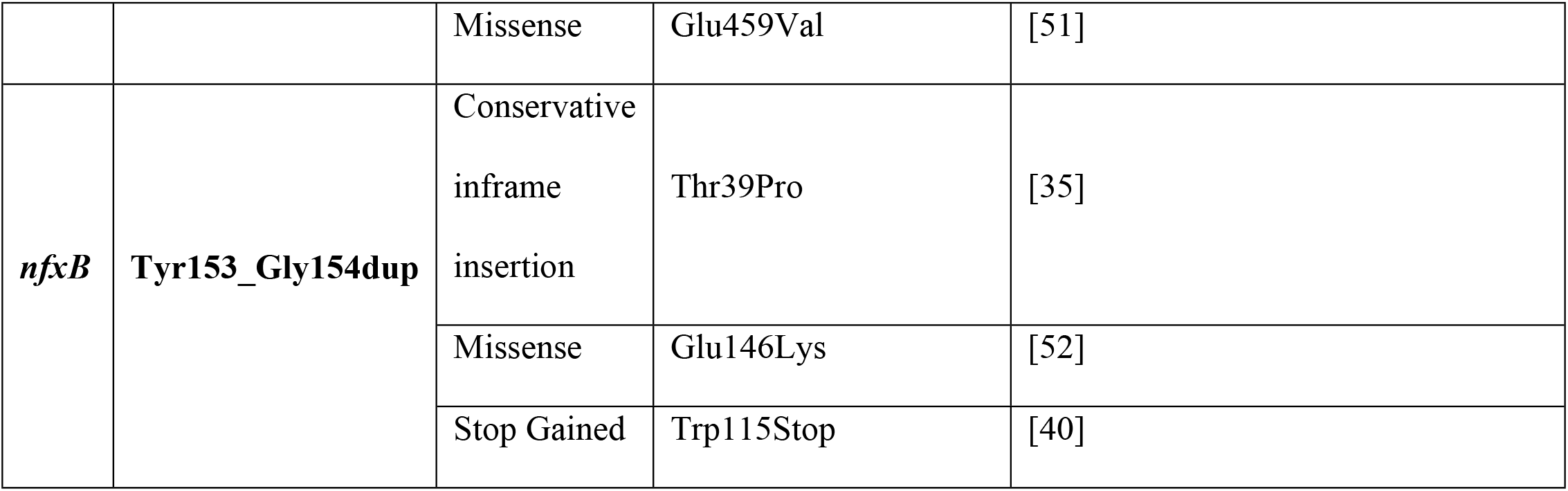
Comparative analysis of mutations identified in the experimentally evolved ciprofloxacin-resistant (CIP-R) *Pseudomonas aeruginosa* strain with previously reported resistance-associated variants described in clinical isolates.

The key factors associated with resistance were remarkably similar in comparison with clinical ciprofloxacin-resistant isolates. The *gyrA* Thr83Ile mutation observed in the CIP-R strain was found in 86.38% of 492 clinical isolates confirming its role in the development of resistance. However, mutations impacting secondary targets were variable. Most of the clinical isolates had *parC* mutations, e.g., Ser87Leu (70.93%) or Ser87Trp, whereas the CIP-R strain exhibited a Thr177Asn substitution. The resistant strain carried the Glu459Lys *parE* mutation, but clinical isolates commonly exhibited Glu459Asp, Ala473Val or Ser457Gly substitutions. Together, our data imply that the core processes of ciprofloxacin resistance are conserved, while the particular mutational patterns may change between laboratory-evolved and clinical strains.

## 4. Discussion

Our study of a laboratory-produced ciprofloxacin resistant strain of *Pseudomonas aeruginosa* indicates that ciprofloxacin resistance is more complex than just mutations in *gyrA*, *parC* and *parE*. Additional mutations were found in *pilA*, *tadB*, *psdR*, *dppA3* variants, *NP446_RS32055* (TRAP transporter permease), and the prophage-associated hypothetical gene *NP446_RS24255* during in vitro evolution under increasing ciprofloxacin pressure. These changes likely reflect broader bacterial adaptation and may contribute to the metabolic trade-offs observed in the resistant strain. The known role of *nfxB* is to repress efflux pumps, but loss of function may lead to overexpression of efflux pumps [20,53]. This is considered to remove excess ciprofloxacin from the bacterial cell. We believe this is indirectly related to multiple fitness trade-offs during high-level ciprofloxacin resistance. Hence, *nfxB* seems to be indirectly associated with such phenotypes. The exact mechanism is not clear from our study and it needs dedicated investigation.

The *gyrA* Thr83Ile substitution is among the most common mutations conferring ciprofloxacin resistance in *P. aeruginosa* and reduces binding affinity of ciprofloxacin [54,55]. The *gyrA* Thr83Ile mutation was conserved in both the laboratory-evolved CIP-R strain and in 86.38% of clinical isolates, suggesting that exposure to ciprofloxacin invariably selects this key genetic determinant of resistance. In addition to Thr83Ile, *gyrA* Asp87Asn mutation was found in 52.85% of the clinical isolates. The laboratory-evolved ciprofloxacin-resistant strain acquired a *parC* Thr177Asn substitution, whereas Ser87Leu was the dominant *parC* variant in clinical isolates (70.93%). On the other hand, *parE* Glu459Lys mutation identified in the CIP-R strain was rarely detected among the clinical isolates. These results indicate that mutational pathways are specific to laboratory evolution and clinical settings and that secondary mutations in topoisomerase IV further increase ciprofloxacin resistance by relieving antibiotic-induced DNA damage. The high-level fluoroquinolone resistance has been reported to involve the stepwise alterations in the genes of DNA gyrase and topoisomerase IV [56]. The exceptionally high MIC of the CIP-R strain correlated with the presence of several target-site mutations.

The CIP-R strain exhibited drastically reduced swimming and swarming, but twitching motility was unaffected. Although we observed mutations in *pilA* and *tadB*, which are involved in pilus formation and surface-associated motility. Type IV pili are essential for adhesion, biofilm maturation, and coordinated surface motility of *P. aeruginosa* [57]. Thus, the reduction in biofilm observed in our results may be a consequence of mutations in these two genes. However, the reduction in swimming and swarming seems to be more related to fitness trade-offs and might be indirectly associated with *nfxB* mutations. In one study, a mutation in *nfxB* led to a reduction in motility [58]. Similarly, the reduction of EPS is also related to lot of physiological response. The resistant strain may have distributed its metabolic resources into resistance mechanisms rather than biofilm formation. SEM analysis revealed weaker surface association, less clustering and smaller microcolonies in the resistant strain. CLSM further confirmed that the CIP-R strain had a higher cell viability against the exposure of ciprofloxacin, while the parental strain had more cellular damage. The collateral sensitivity and resistance patterns observed in the resistant strain shows the evolutionary impact of adaptation to antibiotics. The evolved strain exhibited cross-resistance to levofloxacin, cefepime and meropenem but increased susceptibility to piperacillin– tazobactam and aztreonam. Resistance to levofloxacin is likely due to the common targets of fluoroquinolones, but altered responses to other antibiotics may be due to more general physiological responses such as efflux activity or membrane changes. These results imply that the evolution of resistance can simultaneously create new vulnerabilities that can be exploited therapeutically.

Overall, our study reveals that extended ciprofloxacin treatment results in substantial genomic, physiological and phenotypic adaptation of *Pseudomonas aeruginosa*. Progressive accumulation of mutations in *gyrA*, *parC* and *nfxB* was linked with stable high-level resistance, reduced motility, weak biofilm formation, slowed growth, changed surface attachment and collateral changes in antimicrobial sensitivity. The high prevalence of *gyrA* mutations supports its utility as a potential biomarker of ciprofloxacin resistance, whereas mutations in *parC* and *parE* may vary under in vivo and in vitro conditions. We believe that all the physiological changes were associated with *nfxB* mutations. For instance, the evolved strain shows a reduction in biofilm formation despite the absence of mutations in key biofilm-associated genes. Based on our collateral sensitivity data, the CIP-R strain can be treated with beta-lactam antibiotics, although a beta-lactamase inhibitor still needs to be added. The potential efficacy of bacteriophage therapy should also be investigated, as the *nfxB* mutation appears to influence multiple bacterial phenotypes.

## 5. Acknowledgments

The authors would like to thank VIT, Vellore for providing essential research facilities for conducting this work. The authors also thank the Central Research Facility staff for their assistance with scanning electron microscopy (SEM) and confocal microscopy studies. In addition, the authors acknowledge BioRender.com for the preparation of scientific illustrations and figures.

## 6. Author Contributions

AS: Original draft writing, formal analysis, figures, review & editing, methodology, and investigation. RR: Project management, funding procurement, supervision, resources, final review, and editing.

## 7. Data Availability

Whole-genome sequencing data and assemblies of *Pseudomonas aeruginosa* CIP-R have been submitted in the NCBI GenBank under the BioProject ID PRJNA1475186 with the accession number JBZYDH000000000.

